# Mapping antibiotic resistance determinants in oral streptococci

**DOI:** 10.64898/2026.02.06.700101

**Authors:** Kazi Shefaul Mulk Shawrob, Andreas Solberg Sagen, Tracy Munthali Lunde, Biramitha Sribasgaran, Saba Ghafoor, Mohammed Al-Haroni, Rebecca Ashley Gladstone, Gabriela Salvadori, Roger Junges

## Abstract

**Background:** Antibiotic resistance is a global priority in healthcare. Leveraging thousands of whole-genome sequences, here we reveal the core resistance determinants of oral streptococci, focusing on assessing pattern variability and gene exchange across commensals and pathogens.

**Methods:** Genomic information was obtained from the National Center for Biotechnology Information. Determinants of antibiotic resistance were identified using AMRFinderPlus and CARD. ICEscreen was employed for calling of integrative and conjugative elements. Variability and recombination in penicillin-binding protein sequences were assessed with MMseqs2 and fastGEAR.

**Results:** A total of 2,087 genomes from 15 species were included with members from the mitis, mutans, anginosus, salivarius, and bovis groups. We observed 3,576 hits from 55 unique resistance genes conferring resistance to 11 antibiotic classes. The species with most resistance determinants per genome were identified as *Streptococcus mitis* (2.7), *Streptococcus oralis* (2.5), *Streptococcus parasanguinis* (1.9), *Streptococcus gallolyticus* (1.3), and *Streptococcus anginosus* (1.2). The two latter species also presented the most diverse composition of determinants. Over 1,800 integrative and conjugative elements were shown across all genomes, with nearly 18% of them carrying at least one antibiotic resistance gene. Penicillin-binding protein variation analyses showed a high diversity in the mitis group. Even though *S. mitis* and *S. oralis* composed less than 4% of the genomes included in the analyses, they were recognized as sources of DNA for over a third of recombination events in *pbp1a* and nearly half for both *pbp2b* and *pbp2x* in resistant isolates of *Streptococcus pneumoniae*.

**Conclusions:** We show that tetracycline and macrolide resistance were highly abundant and tightly connected to integrative and conjugative elements. Further, recent recombination data show frequent genetic exchange from oral streptococci to beta-lactam-resistant *S. pneumoniae*. Finally, assessing the dynamics of genetic exchange across species is central for the development of strategies to mitigate the impact of antibiotic resistance.

## Background

The antimicrobial resistance (AMR) crisis is one of the largest threats to global healthcare, as it not only undermines the effectiveness of available antibiotics for treating infections but also endangers many modern medical procedures that rely on antimicrobials. While significant resources are directed toward surveilling the emergence and spread of AMR determinants in microbes worldwide, most efforts are centered on species with high pathogenic potential. In turn, the emerging view suggests that, in addition to pathogens, monitoring such determinants in commensal species can enhance and assist existing surveillance systems worldwide (Goytia and Wadsworth 2022; Vihta et al. 2021). In fact, most AMR genes could be present in commensal species, which may act as a reservoir creating a diverse pool of genes available, thus highlighting the need to investigate carriage and the dynamics of exchange in this context as well (Anderson et al. 2023; Hendriksen et al. 2019; Nji et al. 2021).

The genus *Streptococcus* is one of the most abundant groups of bacteria in the oral cavity. While generally associated with a healthy state, some species are linked to a variety of oral diseases such as caries and mucosal infections (for review see (Bloch et al. 2024)). Commensal streptococci have also been often isolated from sterile sites in cases of severe bacteremia and endocarditis, with an increased occurrence in immunocompromised individuals (Colomba et al. 2023; Marsland et al. 2024; Plainvert et al. 2023; Ronning et al. 2024; Shelburne et al. 2014). While a recent study focused on population genomics of *Streptococcus mitis* in bloodstream infections in the UK and Ireland showed that infective endocarditis has not been primarily caused by hypervirulent or antimicrobial-resistant lineages over the years (Kalizang’oma et al. 2024), other reports indicate a higher rate of oral streptococci non-susceptible to antibiotics recently, with a particular concern for resistance to macrolides, cephalosporins, and carbapenems (Colomba et al. 2023; Marsland et al. 2024; Plainvert et al. 2023; Ronning et al. 2024; Sarangi et al. 2024; van Prehn et al. 2019). In addition, oral streptococci are genetically closely related to *Streptococcus pneumoniae*, a major human pathogen, sharing habitat in the upper airways.

The spread of AMR in bacterial populations relies on horizontal gene transfer (HGT), which often occurs via mobile genetic elements (MGEs) such as integrative and conjugative elements (ICEs). These elements possess the genetic machinery necessary to self-assemble as well as the cargo, which often expresses antibiotic resistance cassettes, virulence factors, and stress response systems. They are considered major drivers for the spread of antibiotic resistance in addition to being central in promoting genomic plasticity for adaptation to environmental stressors (Botelho and Schulenburg 2021; Colombi et al. 2024; Wozniak and Waldor 2010). HGT via natural transformation is also of particular interest in streptococci, as beta-lactam resistance is mediated by mutations in the genes coding for the penicillin-binding proteins (*pbp*s), essential for building the streptococcal cell wall (for review see (Hakenbeck et al. 2012)). The mosaic of alterations across *pbp*s, particularly *pbp1a, pbp2b*, and *pbp2x*, can act in synergy conferring allelic resistance to not only penicillin, but also cephalosporins and carbapenems (Chewapreecha et al. 2014; Hakenbeck et al. 2012). Combination of *pbp* mutations leading to resistance phenotypes has been shown to arise primarily from recombination events and not *de novo* mutations (Nishimoto et al. 2022).

Nonpneumococcal species of the mitis group have traditionally been the focus of studies looking into antibiotic resistance genes (ARGs) and alleles such as mutated *pbp*s (Jensen et al. 2015; Kalizang’oma et al. 2021; van der Linden et al. 2017), nevertheless, as genomic data for non-mitis group streptococcal species is increasingly available, understanding their landscape of resistance is likely to benefit the field. To address these knowledge gaps, we leveraged over 2,000 oral streptococcal genomes to characterize the landscape of antibiotic resistance determinants across species. Here we present a strategy for assessment of antibiotic resistance in oral streptococcal genomes and show that these genetic determinants are pervasive in oral streptococci. We further show that macrolide and tetracycline resistance are widespread and often located in mobile genetic elements, facilitating recombination across species. Finally, by assessing recombination events in the *pbp*s, we observed that oral streptococci were traced as source of DNA for over 40% of recombination events in the *Streptococcus pneumoniae*, despite encompassing about 15% of the genomes included in the analyses.

## Materials and Methods

### Genomic data

The study was conducted using publicly available genomes of 15 streptococcal species (Table S1). Reference genome sequences for these species were retrieved from the NCBI RefSeq database on April 7, 2025. For *pbp* genetic analyses, 10,572 *S. pneumoniae* RefSeq genomes were included additionally. Corresponding metadata files were processed to extract relevant genomic information, including assembly statistics, taxonomic classifications, and sequence identifiers.

### Antimicrobial resistance gene identification

Antibiotic resistance gene (ARG) identification was performed with an approach utilizing the databases of the National Center for Biotechnology Information (NCBI) via AMR Finder Plus (Feldgarden et al. 2021) and the Comprehensive Antibiotic Resistance Database (CARD) (Alcock et al. 2023) via Abricate (Seemann https://github.com/tseemann/abricate). For AMR Finder Plus genes, virulence factors, and stress resistance genes were identified using AMRFinderPlus (version 4.0.19) with the default database updated to the most recent version (2024-12-18.1). Additional AMR gene detection was performed utilizing Abricate (version 1.0.1) with the Comprehensive Antibiotic Resistance Database (CARD, 2025-Jan-14). We defined the multidrug resistance (MDR) score per genome as the number of distinct antibiotic classes represented by its resistance genes as a continuous variable and not a binary classification (Pires et al. 2021; 2022; Shawrob et al. 2025). Categorical MDR was called if the score was 3 or higher, indicating resistance to three distinct antibiotic classes. Further details on dataset curation and harmonization are available in the supplementary material.

### Integrative and conjugative elements (ICE) identification and coupling with ARG data

To identify mobile genetic elements in the genomes, ICEscreen was selected due to its particular focus on detection in Firmicutes (Lao et al. 2022). All genomes were analyzed using ICEscreen v1.3.3 with the phylum taxonomic group set to bacillota. In cases where the detection capability of ICEscreen is in doubt, or extra validation was needed, BLASTn and tBLASTn were used with a query sequence and a custom database created from whole genome, or genome-subsets. Following identification of ICEs, and ARGs, the two datasets were analyzed by identifying ICEs and ARGs co-located by an ARG existing in or near the ICE. To determine whether these ARGs are associated with ICEs, a custom script written with BioPython v1.18 was created with an approach similar to previous reports (Dechene-Tempier et al. 2025; Johansson et al. 2025). Briefly, the algorithm checked if any identified ARGs were located in between the left and rightmost signature protein (SPs). Because ICEscreen does not define the exact ICE boundaries, ARGs may also occur just outside these SPs. To capture these, the script additionally examined a 20 kbp region flanking the SP. When an integrase was identified as either the leftmost or rightmost SP, that boundary was considered definitive, and the flanking search was performed on the opposite side.

### Penicillin-Binding Proteins (PBPs) analyses

The gene and protein sequences of the three main penicillin-binding proteins (PBP), namely PBP1a, PBP2b, and PBP2x were extracted from each isolate and separated by PBP type to create individual datasets for analysis. Protein sequences were clustered individually using MMseqs2 (Steinegger and Soding 2017). For each dataset, sequences were converted into MMseqs2 databases and clustered with graph-based mode, which provides order-independent results. For recombination analyses, *pbp1a, pbp2b* and *pbp2x* sequences from streptococcal species were aligned with MUSCLE using the super5 algorithm on nucleotide input with four iterations and 400 refinement iterations on the nucleotide input. The resulting alignments were used as input for fastGEAR (Mostowy et al. 2017) to detect horizontal gene transfer (HGT) events. Briefly, fastGEAR uses BAPS (Bayesian Analysis of Population Structure) (Cheng et al. 2013) to first generate sequence clusters for each gene. These clusters were subsequently utilized to identify recombination lineages by merging clusters that shared a common ancestry in at least 50 percent of sites via a hidden Markov model. FastGEAR then scanned each lineage with an HMM to detect recent recombinant fragments. Each fragment was assigned to the donor lineage with the highest probability at that position. A Bayes factor value above one was used to accept a recent recombination event (logBF > 0) and each fragment was assigned to the donor lineage with the highest probability. Further details are described in the supplementary material. All outputs were imported into R and processed with Tidyverse. Recombination patterns were visualized with ggplot2. Resistance phenotype in *S. pneumoniae* was predicted utilizing an *in silico* pipeline previously established (Li et al. 2017; Li et al. 2016).

## Results

### Genomic data and strategy for detection of antimicrobial resistance genes

Species of oral streptococci included in this study were based on phylogenetic clustering (Abranches et al. 2018) and presented representatives from the mitis, salivarius, bovis, mutans, and anginosus groups (Table S1). Data from Refseq, which is maintained by NCBI and offers a curated, non-redundant database, were utilized in the analyses (Pruitt et al. 2007). Most of the genomes available were from the mitis group (n=922), followed by mutans (n=446), salivarius (n=318), and anginosus groups (n=348). Sample origin was identified as the oral cavity (725), followed by the gut and fecal sources (245), and blood or heart-related infections (73). For a comprehensive approach, we defined the strategy as a combination of AMR Finder Plus and CARD ensuring a representative and complete selection of ARGs (Supplementary material and Table S2).

### Panorama of ARG presence across oral streptococcal species

The 2,087 genomes were screened for ARGs providing a total count of 3,576 hits for 55 unique genes (Table S3). Macrolides and tetracyclines were the two antibiotic classes that were most frequently found with 1,296 and 844 counts, respectively. At the individual ARG counts, the tetracycline resistance gene *tet(M)* was the most common with 327 counts, being present in 15.7% of all genomes surveyed and across all species with the exception of *S. sobrinus* and *S. downei* from the mutans group. The macrolide resistance genes *erm(B)* and the complex *mef(A)*/*msr(D)* were also nearly ubiquitously identified across all the species (Fig 1A). In contrast, the genes *tetA(46)* and *tetB(46)* were only identified in *S. parasanguinis*. The potential resistance genes for fluoroquinolone resistance *patAB* and *pmrA* were exclusively found in the mitis group (Fig 1B). The two species that presented most diversity of genetic determinants were *S. anginosus* (30 unique genes) and *S. gallolyticus* (27 unique genes). Conversely, only 12 ARG counts were identified in 446 genomes from the mutans group. Multidrug resistance was present in 14.6% of the analyzed genomes, with most common species being *S. mitis* (56.6%), *S. oralis* (40.5%), and *S. gallolyticus* (26.4%) (Fig S1). We further attributed an MDR score to all isolates that counts resistance to antibiotic classes cumulatively (Fig 2A). As such, it was observed that the average score for these genomes was 1.16 (Fig 2B), indicating a little over 1 ARG per genome surveyed. Species with the highest scores were *S. mitis* (2.7), *S. oralis* (2.5), *S. parasanguinis* (1.9) from the mitis groups, followed by *S gallolyticus* (1.3) and *S. anginosus* (1.2).

**Fig 1.**
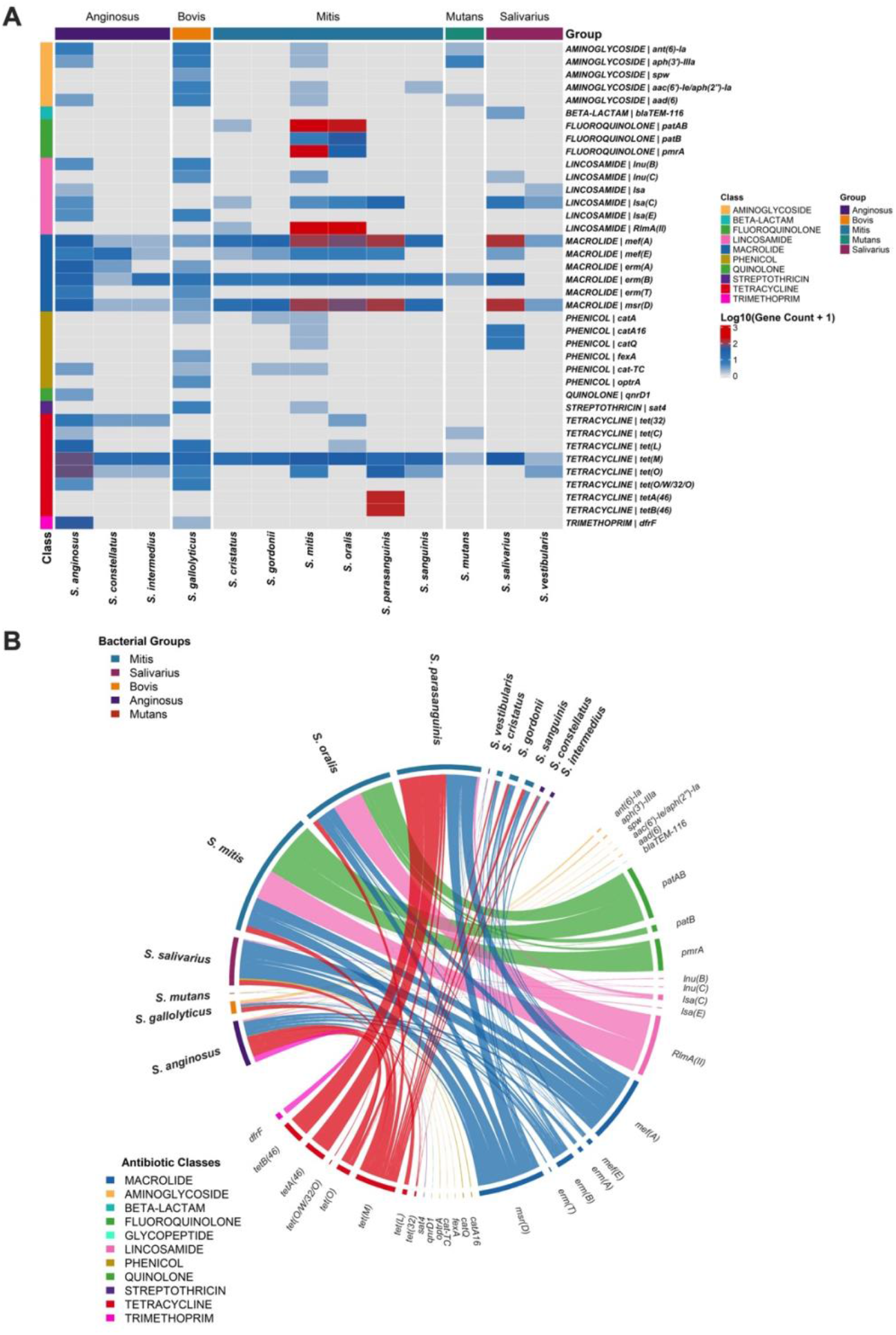
Antibiotic resistance gene distribution across oral streptococci included in this study. **(A**) Heatmap showing the distribution of ARGs across anginosus, bovis, mitis, mutans and salivarius groups. Each row represents a specific gene detected by AMRFinderPlus and/or CARD. Genes are grouped by antibiotic class, and each column represents a species. Colors intensity shows gene count in log10 scale. Genes with a count lower than two were excluded. **(B)** Chord diagram showing the associations between oral streptococci and resistance genes. Bacterial species were categorized into five phylogenetic groups and individual ARGs are color-coded by their associated antibiotic class. Bandwidth represents the frequency and association of the ARGs within bacterial group. Genes with a count lower than two were excluded.

**Fig 2.**
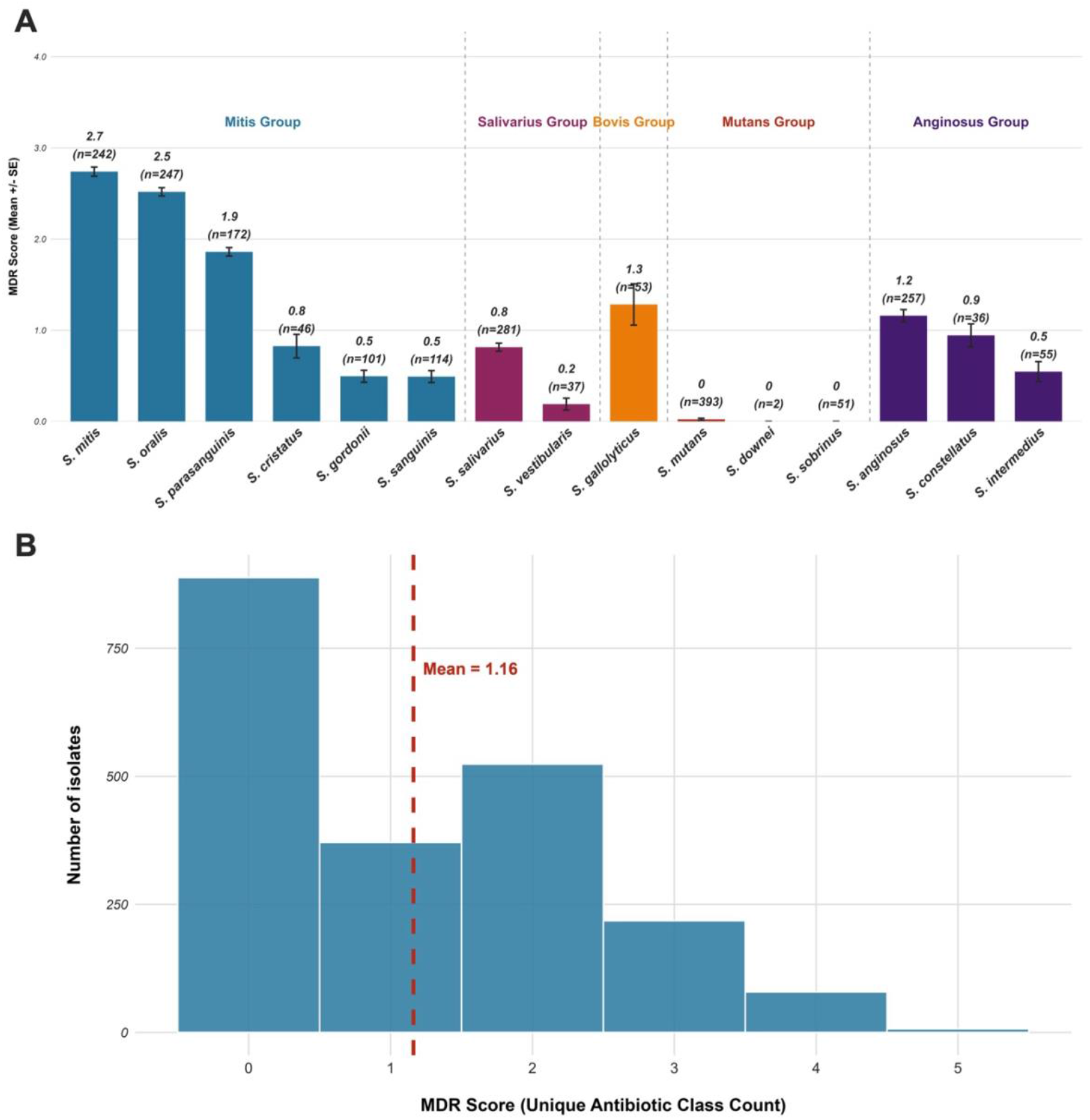
Multidrug resistance score across oral streptococci groups included in the study. **(A)** Antimicrobial resistance score of streptococcal species groups. Each bar shows the mean MDR scores, representing the count of unique antibiotic classes to which a species is resistant to for individual species. Error bars indicate the standard error of the mean. Number of genomes for each species are listed above each bar. **(B)** Distribution of multidrug resistance scores among all oral streptococcal genomes. The histogram shows the frequency of MDR scores across the total population of isolates studied. The scores range from 0 to 5 unique antibiotic classes, with a calculated overall mean MDR score.

### Linkage between integrative and conjugative elements (ICEs) and resistance genes

Given the major role of ICEs in promoting genomic plasticity and lateral exchange of cargo, we sought to understand the current profile of antimicrobial resistance associated with these elements in oral streptococci. Utilizing ICEscreen (Lao et al. 2022), we identified a total of 1,829 putative complete ICEs, resulting in 0.87 ICEs per genome surveilled. Additionally, 637 partial ICEs (0.25 per genome) and 295 conjugation modules (0.19 per genome) were identified. In total, ~63% of all genomes surveilled contained at least one putative ICE or ICE-like element, with 50% of genomes containing at least one complete ICE (Table S4). *S. anginosus* harbored most ICEs both in absolute and relative terms (2.25 putative complete ICEs per genome), while *S. gallolyticus, S. constellatus, S. intermedius* and *S. sanguinis* also carried more than one complete ICE per genome on average. The distribution of ARGs and ICEs across species indicates variations across groups (Fig 3A). The most frequently detected ICE subfamily was Tn*916*, and the sizes of these putative elements largely aligned with the expected size range for this family, though some variation was observed. Tn*5252* was the second most common subfamily identified (Fig 3B). Among the complete ICEs identified, 17.9% contained an ARG, while 12.4% of partial ICEs and 21.4% of conjugation modules were associated with an ARG. *S. cristatus* carried the most ARGs per complete ICE (37.1%) and had the highest frequency of ICEs, carrying 2 or more ARGs at 20% of the total number of ARGs. *S. mitis* was found to harbor the second highest frequency of ARGs associated with a complete ICE at 29.4% with 14.7% containing two or more ARGs. Tetracycline and macrolide resistance genes were the most frequently associated with ICEs, particularly *tet(M)*, and *erm(B)* (Fig S2).

**Fig 3.**
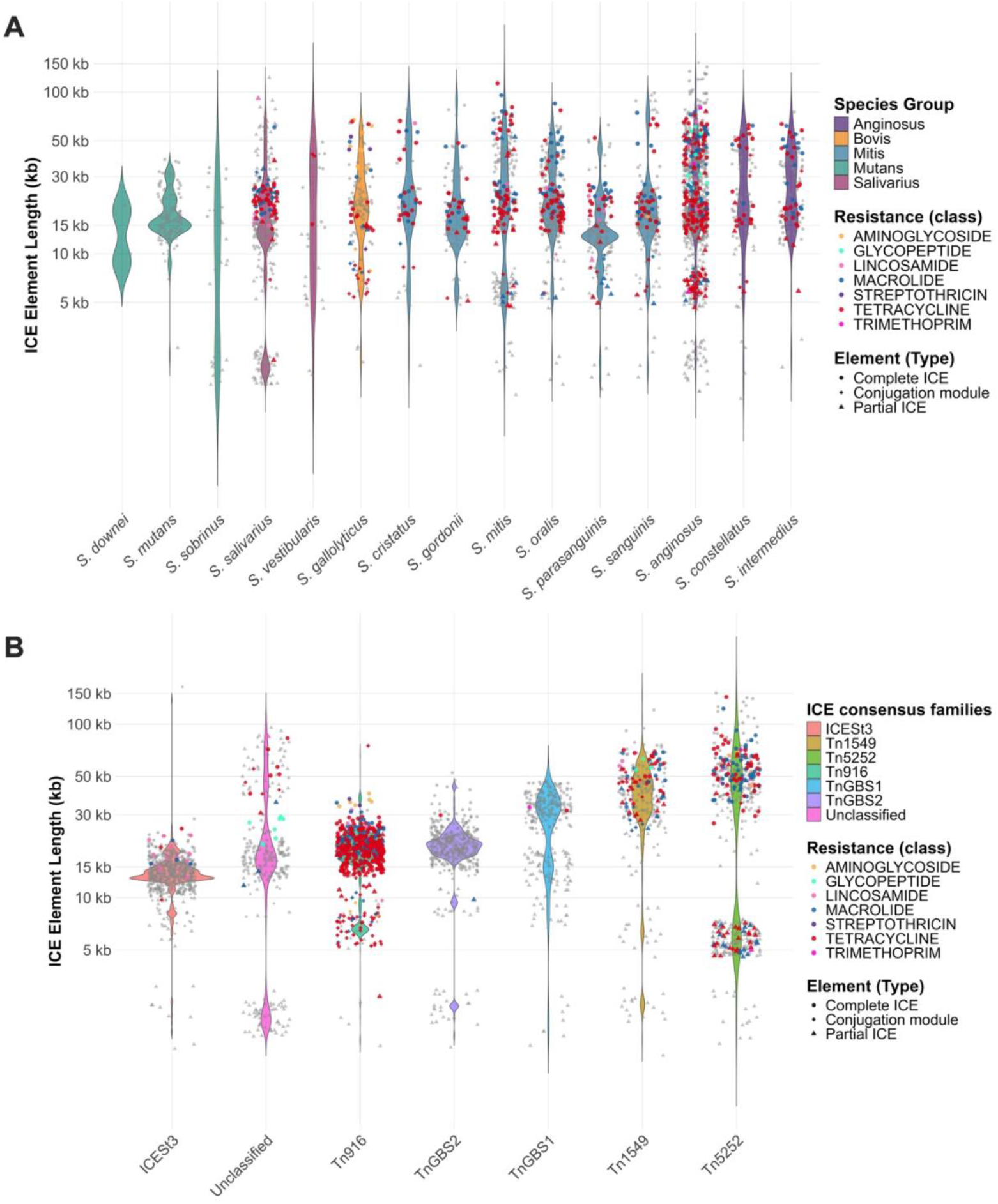
Complete ICE, partial ICE and conjugation module distribution across the genomes surveilled in the study.**(A)** Size distribution of each identified integrative and conjugative element across different species. Each dot represents an element with an identified ARG causing resistance and classified as either ARG-inside or ARG-proximal depending on if the ARG was identified within the boundaries of the ICEs SPs or outside but within 20kbps from the non-integrase edge of the element. **(B)** Size distribution of each identified integrative and conjugative element across different subfamilies. Each dot represents an element with an identified ARG causing resistance and classified as either ARG-inside or ARG-proximal depending on if the ARG was identified within the boundaries of the ICEs signature proteins or outside but within 20kbps from the non-integrase edge of the element.

### Diversity in the penicillin-binding protein genes and recombination across species

As mutations in the *pbp*s coordinate resistance to beta-lactams in streptococci, we aimed to reveal their diversity patterns. Firstly, we extracted their sequences for the genomes included in this study, as well as 10,572 sequences of *S. pneumoniae*. We then clustered all protein sequences at a 95% threshold to assess diversity (Fig 4). Species from the mitis group, particularly *S. pneumoniae, S. mitis*, and *S. oralis* presented significant divergencies in *pbp* profiling compared to the other species. Moreover, significant overlap of species across clusters was observed. While in *S. pneumoniae* and *S. oralis* a cluster with the majority of sequences was observed for all *pbp*s, in *S. mitis* it was more challenging to define a most common type. The discrepancies in number of sequences compared to the number of clusters formed in *S. pneumoniae* and *S. mitis* further emphasizes the diversity in the latter. Predicted phenotypic resistance in *S. pneumoniae* showed that while the major clusters showed susceptibility, resistance tended to be concentrated in specific clusters. This was particularly relevant for PBP1a, where nearly all highly resistance strains were clustered together (Table S5).

**Fig 4.**
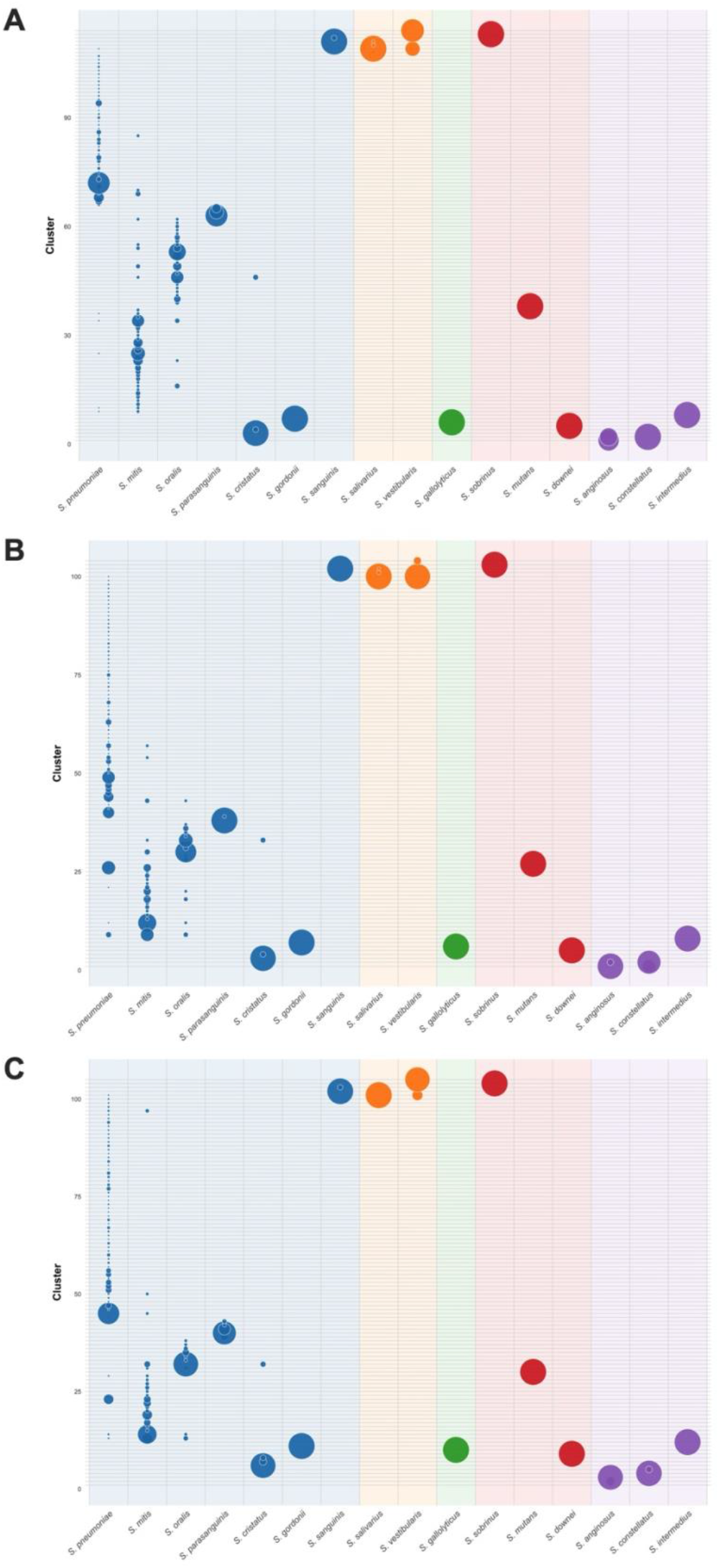
Distribution of PBP protein clusters across all oral streptococci groups and *S. pneumoniae*. Sequence diversity was analyzed for **(A)** *pbp1a*, **(B)** *pbp2b*, and **(C)** *pbp2x*. MMseqs2 was utilized with a 95% amino acid sequence identity threshold. Species are color-coded by phylogenetic group with bubble size representing the proportion of sequences belonging to a specific cluster within each species.

For the recombination analyses of *pbp* sequences, different patterns were observed in the three *pbps* (Fig 5 and S3). The mitis group, particularly *S. pneumoniae, S. mitis*, and *S. oralis*, showed a high number of recombination events and total DNA transferred. The three aforementioned species served as donors of DNA, with an overrepresentation of *S. mitis* and *S. oralis*, particularly for *pbp2b* and *pbp2x*. For *pbp2x*, over a third of the total DNA exchanged via recombination originated from non-pneumococcal mitis species, despite *S. mitis* and *S. oralis* presenting less than 4% of the genomes available compared to *S. pneumoniae*. High recombination rates were observed in particular for *S. mitis* (Table S6). With data on resistance phenotype prediction, the patterns were similarly observed with commensal streptococci serving as significant donors of DNA to resistant *S. pneumoniae* (Fig 6 and S4). For *pbp2x*, while susceptible pneumococcal genomes showed a recombination rate of less than 9%, resistant phenotypes showed recombination in over 90% of the genomes assessed (Table S7), and in highly resistant strains, the rate was over 99.5%. Overall, oral streptococci composed about 15% of the genomes included in the recombination analyses; however, collectively, they were tracked as a source of DNA for over 40% of recombination events in *S. pneumoniae*. This was mainly driven by *S. mitis* and *S. oralis*, and *pbp2b* and *pbp2x* show high rates of interspecies recombination. While this was still observed in *pbp1a*, intra-species pneumococcal recombination was more often observed in this case.

**Fig 5.**
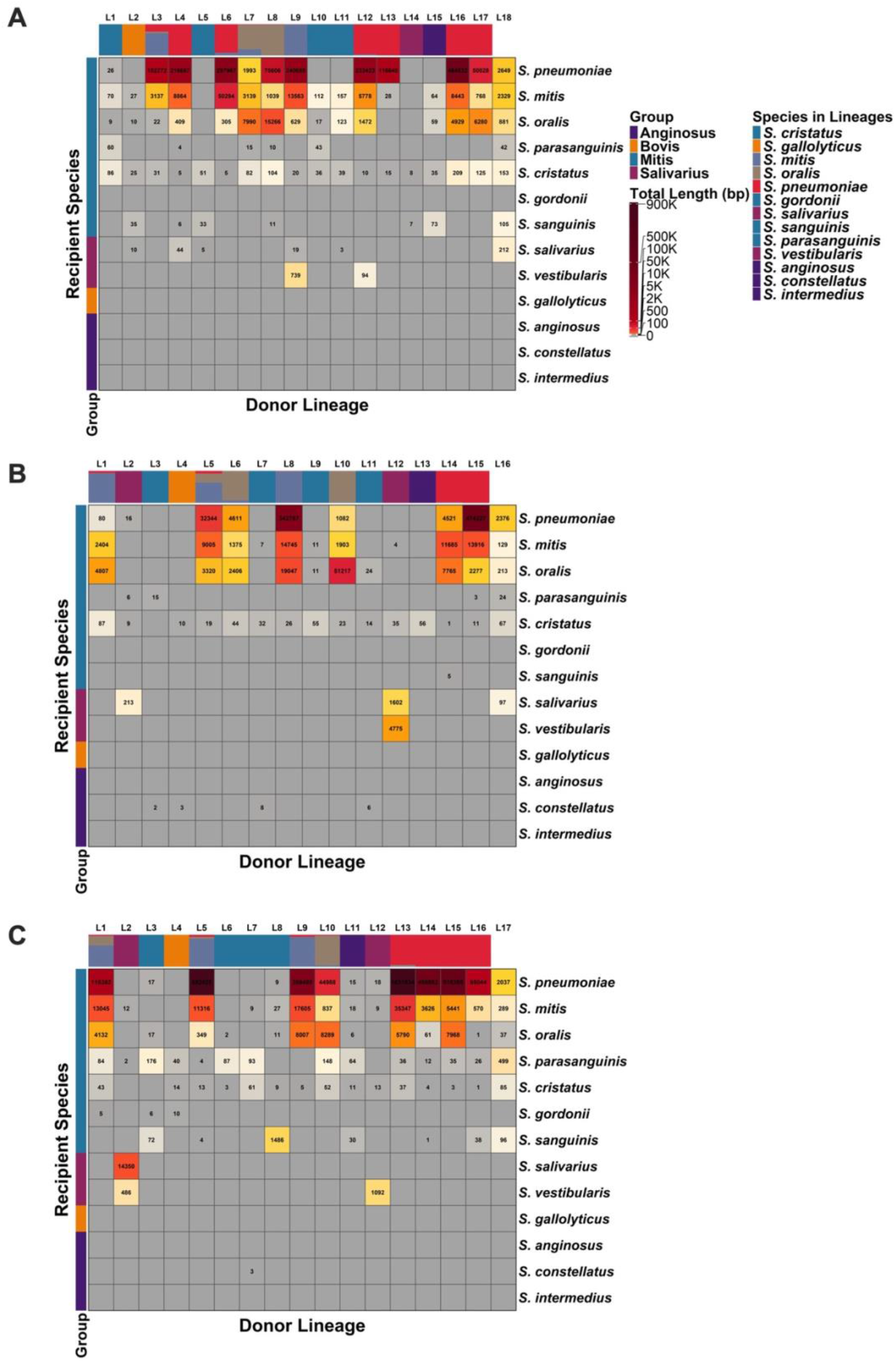
Recent recombination fragments detected by fastGEAR across streptococcal lineages for three penicillin binding protein genes. **(A)** *pbp1a*, **(B)** *pbp2b*, **(C)** *pbp2x*. The heatmap shows transferred DNA fragments from donor lineages (columns) to recipient species (rows). Donor lineages are ordered according to lineage composition shown in the top color bar. Recipient species are ordered according to species group composition shown in the left color bar. Each cell shows the total length of recombination fragments in base pairs adjusted as a heatmap.

**Fig 6.**
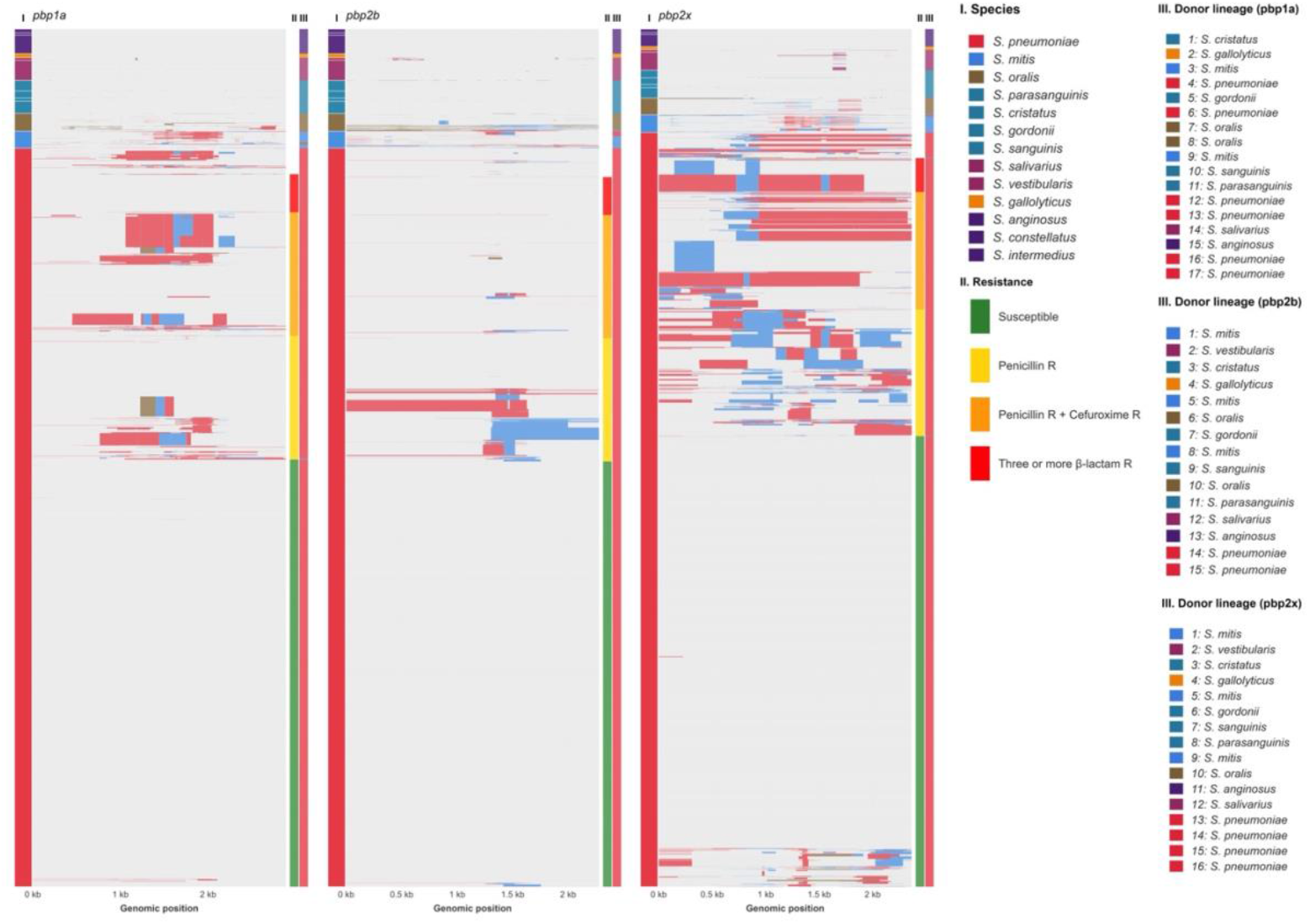
Horizontal gene transfer via recombination events across species in *pbp1a, pbp2b*, and *pbp2x*. For each panel, recipient species are marked on the left side with colors, and donor lineage is marked to the right of the panels. For *S. pneumoniae*, predicted resistance patterns are listed on the right side of each panel and coded by colors. Rows represent individual isolates grouped by species. The x axis shows genomic position in kilobases. Colored blocks across the alignments show recombined fragments detected by fastGEAR. Blocks sharing the same color represent fragments that originate from the same donor lineage based on dominant species detected by fastGEAR recombination lineages. Distinct colors highlight donor contributions from *S. mitis, S. oralis*, and *S. pneumoniae*.

## Discussion

Here we report the core resistome of oral streptococci by leveraging thousands of curated whole-genome sequences. We show that while some of these ARGs are anchored in the chromosome and seem to show little recombination in the population of sequences tested, others are almost exclusively present in ICEs, indicating their high chance of recombination frequency. In addition, we show that the rate of ARG and ICE per genome varies significantly across streptococcal species, with the mutans group exhibiting low counts for both whereas the anginosus group seems to be prolific both in ARG diversity as well as in the presence of uncharacterized ICEs. Furthermore, we investigated diversity and recombination in the *pbps* of these species, with the finding that a high diversity is present particularly in the mitis group. Intra-species and inter-species recent recombination events in this group were observed at a high frequency, and particularly *S. mitis* and *S. oralis* are prolific donors and receivers of genetic fragments from other species.

We report high diversity in the *pbp* genes particularly in the non-pneumococcal species of the mitis group. In other groups of streptococci, these genes were more conserved, often clustering together. In line with our findings, previous studies focused on the mitis group have also reported large heterogeneity in commensal species compared to the pneumococcus (Jensen et al. 2015). We observed that highly resistant strains clustered mostly together in *pbp1a*. Further, cross-species recombination was more often observed for *pbp2b* and *pbp2x*. These findings go in line with previous reports indicating that the *pbp1a* governs high-level resistance in *S. pneumoniae* (D’Aeth et al. 2021; Smith and Klugman 1998). Fewer recombinations were detected in *pbp2b* compared to *pbp1a* and *pbp2x*. We hypothesized that this could have happened due to sequence variations across species resulting in poor alignment. To answer this, we attempted to align only *S. pneumoniae, S. mitis*, and *S. oralis* and observed little differences to the previous data, suggesting that the lack of recombination was not due to poor alignment. It has been reported previously that recombinations in *pbp2b* carry a fitness cost (Albarracin Orio et al. 2011), which could be explanation as to the lower frequency of events observed in our findings. In addition, we corroborate previous findings (Kalizang’oma et al. 2021; van der Linden et al. 2017) and expand on the finding that high variance in *pbp2x* with promiscuous recombination by showing that nearly all resistant strains showed recombination, in comparison less than 10% of the susceptible isolates.

Prolific recombination events were observed in the mitis group compared to other groups, which indicates the mitis group as particularly prone to transformation *in vivo*. As oral streptococci are genetically closely related to *S. pneumoniae*, we have previously shown that tight cell-to-cell contact in biofilms can facilitate genetic exchange of large DNA fragments, likely taking place in the upper airways (Cowley et al. 2018). *S. mitis* and *S. oralis* encompassed less than 5% of pneumococcal genomes and, regardless, they were overrepresented as the source of genetic material. The inclusion of new sequences and public access to a higher quantity of genomes is likely to allow for further research to be developed on the topic. However, it should be highlighted that detection of recombination events is highly dependent on the diversity of genomes included, coupled with the quality of alignment. For instance, in the findings presented here, the mutans group was excluded from recombination analyses due to high divergencies in sequence structure, particularly for *pbp2b*. As such, the data presented here shows bias towards the mitis group of streptococci. We have also explored recombination events in individual groups of streptococci (data not shown), *i*.*e*. anginosus group alone, and we detect more recombination events likely due to better alignment as a product of sequence similarity. While interspecies recombination was our focus due to the potential of transfer to *S. pneumoniae*, it should be noted that future research exploring recombination should be performed with diverse strategies to understand the dynamics of genetic exchange in microbes. Finally, the involvement of other genes in streptococcal antibiotic resistance such as the *murMN* operon has been reported to play a role (Smith and Klugman 2001) and other genes such as *murE* have also been implicated (Todorova et al. 2015).

With 50% of all oral streptococci genomes analyzed containing at least one complete ICE, these elements likely play a significant role in the transfer of ARGs in the oral microbiome. The distribution of such ICEs was uneven across species: members of the mutans group contained considerably fewer ICEs per genome, often less than one, whereas species in the anginosus group exhibited a much higher frequency, averaging 1.75 ICEs per genome, with very few genomes lacking any complete ICE or ICE-like elements. In line with our findings, a recent study revealed a diverse array of chromosomally integrated mobile genetic elements (cciMGEs) in selected oral streptococcal species (Lee et al. 2024). Moreover, their findings included the identification of novel relaxases, highlighting the dynamic nature of these elements in oral streptococci. Overall, ARGs responsible for tetracycline and macrolide resistance were the most observed association with high frequency found in *tet(M)* and *erm(B)* across most species, which goes in line with previous studies assessing presence of ICEs in bacteria from the oral cavity (Wang et al. 2024). As streptococci are the most abundant genus of the oral cavity, these findings can also correlate to the frequent identification of tetracycline and macrolide resistance in saliva samples (Brooks et al. 2022; Vanhout et al. 2024). Noteworthy were the challenges in identifying *mef(A)* and *msr(D)* resistance genes associated with elements via ICEscreen, which have previously been reported to be present in mobile genetic elements (Ambroset et al. 2015; Mingoia et al. 2014). Initially, we hypothesized the issue to be related to genome fragmentation, but upon inspection of the subset of completely assembled genomes, the issue persisted as only two were found to harbor an ICE associated with *mef(A)* and *msr(D)*, both of which being Tn*5252* elements. Employing homology searches with ORF3 and ORF4, which known conserved elements of the macrolide efflux genetic assembly (mega) element harboring *mef(A)* and *msr(D)* in streptococci, 36 and 35 genomes contained fragments of high identity (E≤0.001) out of 37 completely assembled genomes with *mef(A)* and *msr(D)*. Across all genomes irrespective of assembly level, the same ORFs were found in 300 and 299 genomes out of 533 genomes with *mef(A)* and *msr(D)*. The mega element is ∼5.5kb and lacks enzymes required for DNA transposition (Gay and Stephens 2001), as such being non-conjugative on its own. Further, in addition to its presence in composite ICEs as mentioned above, previous reports also indicated its possibility of transfer via natural transformation (Del Grosso et al. 2006; Schroeder and Stephens 2016). Identifying ICEs can be challenging as the backbones can be atypical, the gene order can be scrambled due to recombination, and there is divergence in the core genes. In addition, detection of concatenation across these genetic entities can be difficult in WGS at the contig level. We utilized a fairly strict approach to characterize these elements, similar to previous studies (Johansson et al. 2025; Lao et al. 2022); however, it should be noted that the results here are likely an underestimation of the total of ICEs present given the strictness of our approach. Finally, other MGE components such as integrative and mobilizable elements, phages, and integrons likely contribute significantly to antibiotic resistance transfer and should be further studied.

Despite increases in the last decade, the scarcity of genomes creates challenges for extrapolation and interpretation of the findings at a larger scale. We recently have been able to characterize over 75,000 genomic sequences of *S. pneumoniae*, employing this data to understand temporal and geographical patterns of AMR distribution (Shawrob et al. 2025). While there are over 10,000 curated pneumococcal sequences available at NCBI, for all oral streptococci together there are only 2,000. Surveilling and gathering data on the resistome of commensal species is important to the development of strategies to combat AMR (Anderson et al. 2023; Goytia and Wadsworth 2022; Hendriksen et al. 2019; Nji et al. 2021; Vihta et al. 2021). Altogether, the data presented in this study reinforce the clinical relevance of understanding gene flow between commensals and pathogens, especially in habitats such as upper respiratory tract, where close spatial interactions facilitate natural transformation, which reflects current evidence that interspecies recombination drives the evolution of β-lactam resistance in pneumococci (D’Aeth et al. 2021). In addition, as penicillin is often the first line of treatment for streptococcal infections - primarily from the pathogenic species *S. pneumoniae, Streptococcus pyogenes*, and *Streptococcus agalactiae* - understanding the dynamicity of *pbp* fragments is paramount for diagnostics and surveillance efforts. Nevertheless, it should be noted that while *in silico* prediction of resistance has been available for *S. pneumoniae* (Li et al. 2016), there are no current reliable methods in oral streptococci (Eriksen et al. 2023). Recently, we reported an average MDR score of 0.9 antibiotic resistant determinants per genome in *S. pneumoniae* (Shawrob et al. 2025) and here we show a slightly higher MDR score of 1.2 ARGs in oral streptococci. While the previous analysis did not include intrinsic genes, e.g. *pmrA, patAB*, and *RlmA(II)*, it did include a prediction of *pbp* divergence suggesting resistance. Our study shows the high burden and accumulation of resistance genes/alleles in oral streptococci and their relevance to resistance in major human pathogens. It is important to note, however, that oral streptococcal genomes potentially present a bias risk of resistance overrepresentation compared to more frequently sampled species such as *S. pneumoniae*. Often grouped together as oral or the viridans group of streptococci, our findings call attention for the importance of investigating each group of species for a better understanding of their physiology and ability to act as a reservoir of antibiotic resistance.

## Author contributions

K.S.M. Shawrob, A.S. Sagen contributed to study conception and design, acquisition, analysis and interpretation of data, drafting and critical revision of the manuscript; T.M. Lunde, B. Sribasgaran, S. Ghafoor, M. Al-Haroni contributed to interpretation of data and critical revision of the manuscript; R.A. Gladstone contributed to study design, interpretation of data, and critical revision of the manuscript; G. Salvadori contributed to study conception and design, interpretation of data, and critical revision of the manuscript; R. Junges contributed to study conception and design, analysis and interpretation of data, drafting and critical revision of the manuscript, study supervision. All authors gave their final approval and have agreed to be accountable for all aspects of the work.

## Data availability statement

Data utilized in this study are public and available in the NCBI Genomes Database (https://www.ncbi.nlm.nih.gov/home/genomes/) accessed on Apr 7th, 2025.

## Acknowledgements

Subsets of the analyses described in this paper were performed utilizing the high-performance computing (HPC) clusters owned by the University of Oslo IT Department.

