## Supplementary methods and figures for "Mapping antibiotic resistance determinants in oral streptococci"

### Supplementary material

#### Dataset curation and harmonization

CARD and AMRFinderPlus results were integrated with RefSeq metadata using custom R scripts. Assembly accessions were extracted from AMR output files and matched with bacterial taxonomic information using the readr, stringr, and dplyr packages. Gene names were standardized using a replacement dictionary to harmonize variants (e.g., tet, erm, van gene families) with dplyr::mutate(). Variants of *patA*/*patB* were merged and *mel* genes were standardized to *mef(A)*, while tet, erm, and van gene family variants were standardized under a consistent naming format to ensure accurate gene counting and classification. AMRFinderPlus results were prioritized as the primary source, with CARD predictions added for genes not detected by AMRFinderPlus. Metal resistance genes detected in AMRFinderPlus associated with metal resistance (e.g. *ars*, *mer*, *sil, pco*) were removed from the dataset. Data were grouped by assembly accession and organism name using dplyr functions (group_by(), summarise()), creating a non-redundant combined gene list per genome. Low-quality predictions flagged as INTERNAL_STOP, PARTIAL_CONTIG_ENDX, or PARTIALX were filtered out using stringr pattern matching. Each ARG was mapped to drug classes using a curated classification scheme implemented with dplyr::mutate(). Gene and class information were consolidated per genome. We calculated species-level mean MDR scores by averaging MDR scores across genomes within each species. All data processing utilized Tidyverse ecosystem, with final result exported to excel format using writxl for downstream analysis. To harmonize the results, we employed additional scripts to insert organism names into each file and merge all result files into a single dataset. This combined file was then exported to SPSS, where cross-tabulation was performed to calculate gene frequencies for each bacterial sample. Finally, R Studio was utilized for visualization.

#### Strategy for detection of antimicrobial resistance genes

Typically, *in silico* AMR detection involves the use of search algorithms to screen genomes or protein sequences for known resistance determinants in reference databases (Hendriksen et al. 2019). Thus, screening is dependent both on the strategy utilized by the algorithm as well as the quality and curation of the database (Davies et al. 2023; Papp and Solymosi 2022). As such, with the goal of identifying if there were any major discrepancies for the search in streptococci, prior to assessing the presence of these determinants in all genomes collected, we compared the detection outcomes of a variety of databases utilizing *S. oralis* genomes. AMRFinderPlus (Feldgarden et al. 2021) was utilized in addition to Abricate

(Seemann https://github.com/tseemann/abricate) that automates the search in five databases: CARD (Alcock et al. 2023), ResFinder (Florensa et al. 2022), MEGARes (Bonin et al. 2023), ARG-ANNOT (Gupta et al. 2014), and NCBI AMRFinderPlus (Feldgarden et al. 2019). While nomenclature and gene classification differences ensued as shown in Table S2, the methods utilized presented similar results.

**Penicillin-Binding Proteins (PBPs) analyses**

For each recipient group and species, recent recombination was quantified by total event counts, DNA transfer statistics, including transfer length distributions and DNA transferred per genome. The analyses were performed in *pbp*s in oral streptococci addressed in the study, with additional analyses carried out for *S. pneumoniae*. Minimum inhibitory concentration data for *S. pneumoniae* genomes were predicted utilizing a validated pipeline (Li et al. 2017; Li et al. 2016). Resistance pattern analysis was performed to categorize resistance profiles into groups as shown in tables and figures. MMseqs2 sequence clusters were integrated with isolate (MIC) tables by matching unique accession numbers. This merged dataset subsequently used to analyze the correlation between specific sequence clusters and high resistance levels across all three PBPs. The pneumococcal recombination events and the total amount of DNA transferred (bp) were calculated in R, including the composition of donor lineages in all three PBPs. These data were further integrated with resistance patterns to reveal correlation between recombination events and enhanced beta-lactam resistance. All outputs were imported into R and processed with Tidyverse. Recombination patterns were visualized with ggplot2.

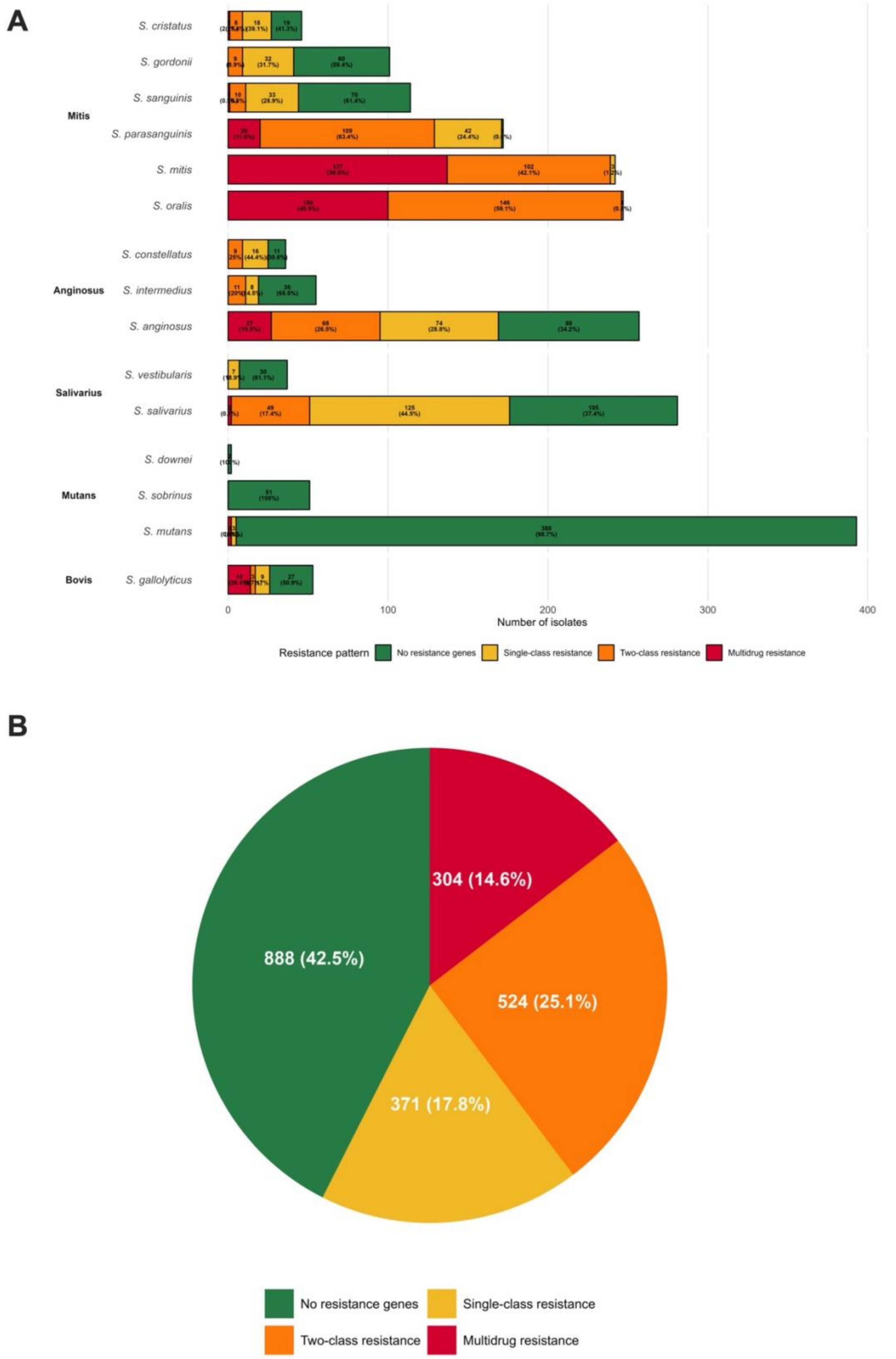


**Fig S1**. Multidrug resistance across oral streptococci. **(A)** Bar plot showing no resistance, resistance to one class, two classes, and multidrug resistance (MDR) (%) per species. **(B)** Resistance to one, two, or three and more antibiotic drug classes in all of the genomes included in the study (n=2,087).

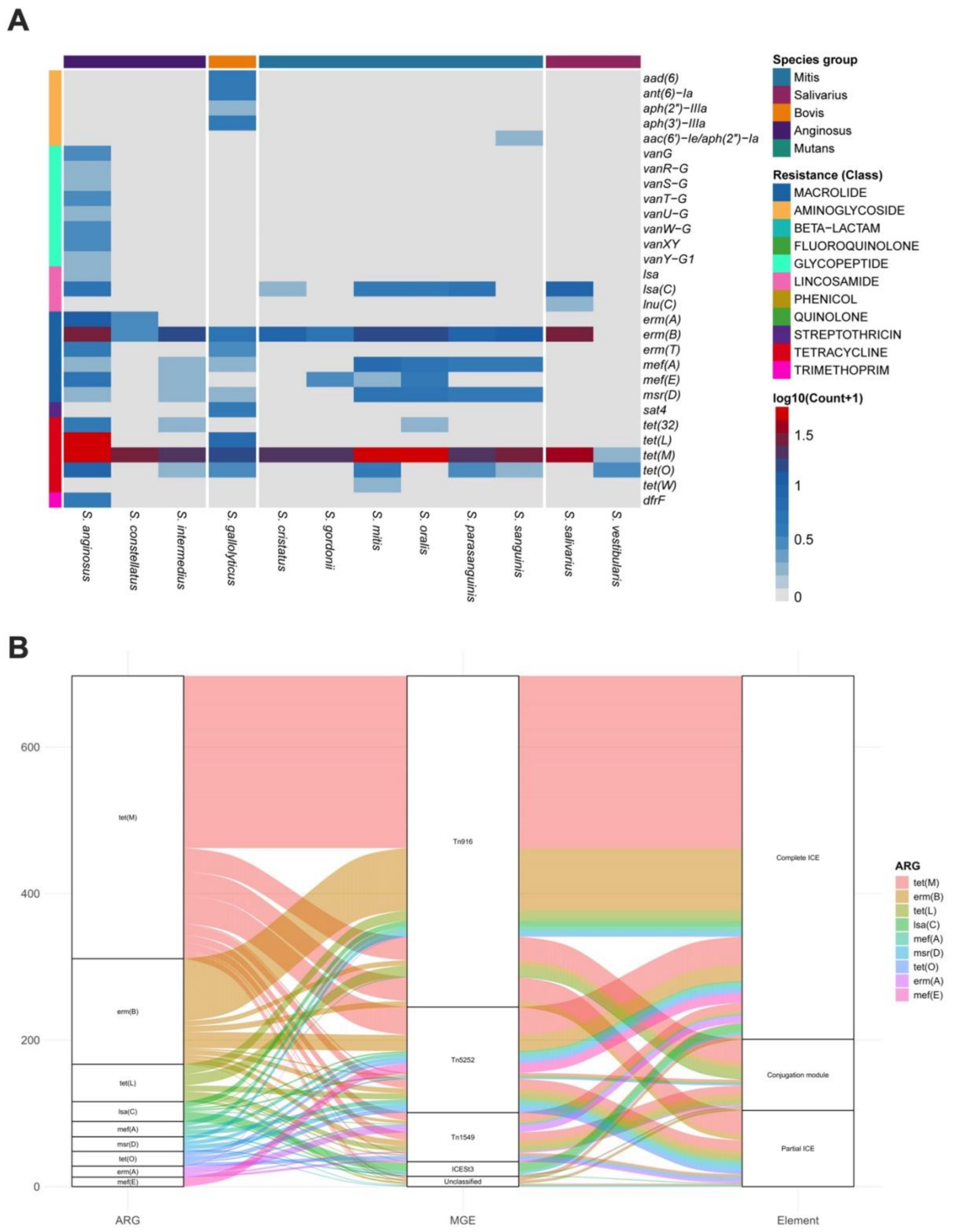


**Fig S2**. ARG distribution in integrative and conjugative(-like) elements. **(A)** Distribution over species. Heatmap shows count of each ARG identified either in or near integrative or conjugative elements across various species. **(B)** Association between the most numerous ARGs (n≥10) and MGE (n≥5) colored by the resistance class of each ARG.

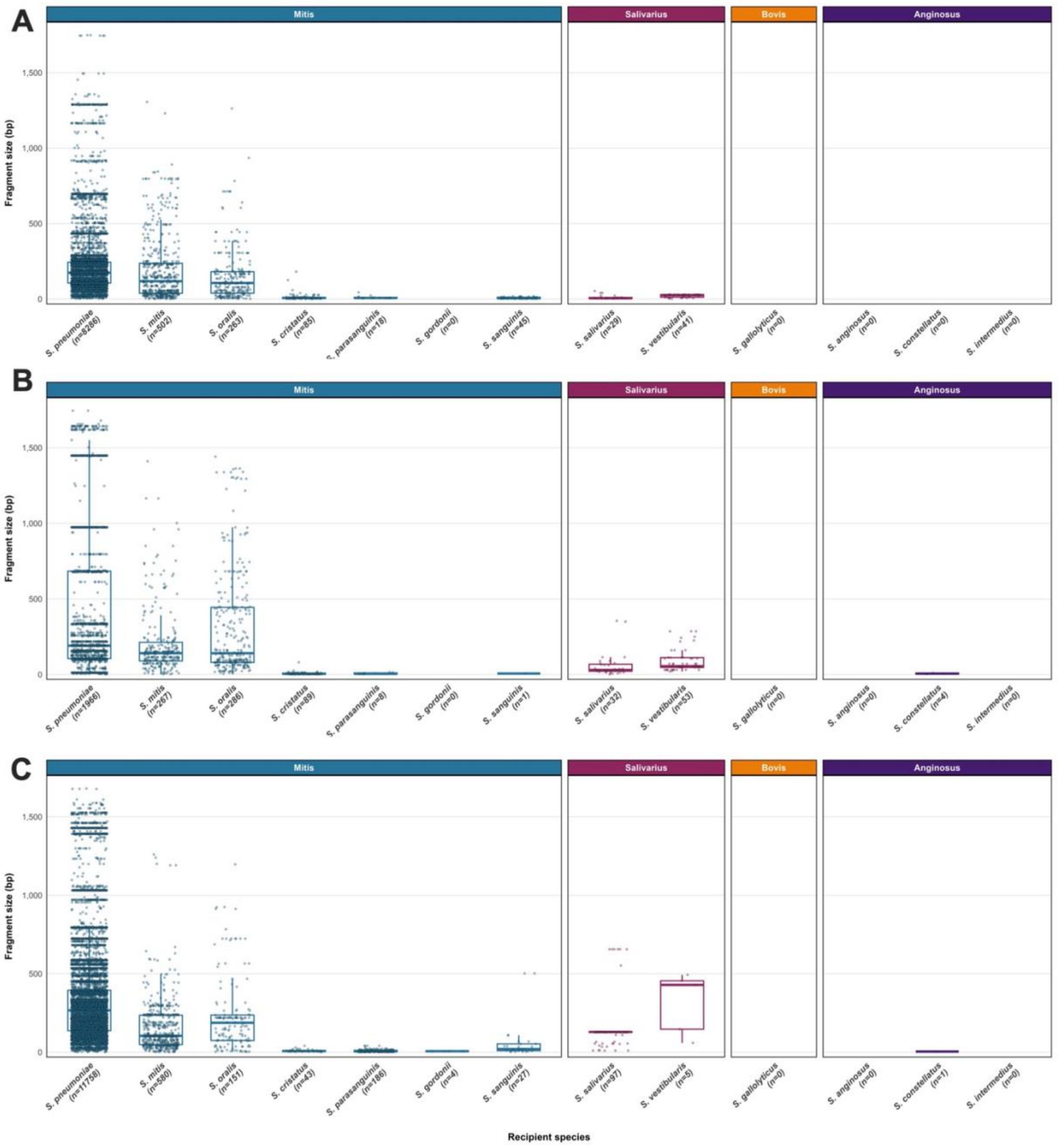


**Fig S3.** Distribution of recombination fragment sizes detected in **(A)** *pbp1a* and **(B)** *pbp2b*, (C) *pbp2x* by species**.** Individual points represent recombination events. Boxplots show median and interquartile range. Numbers below category labels indicate number of events detected in each category.

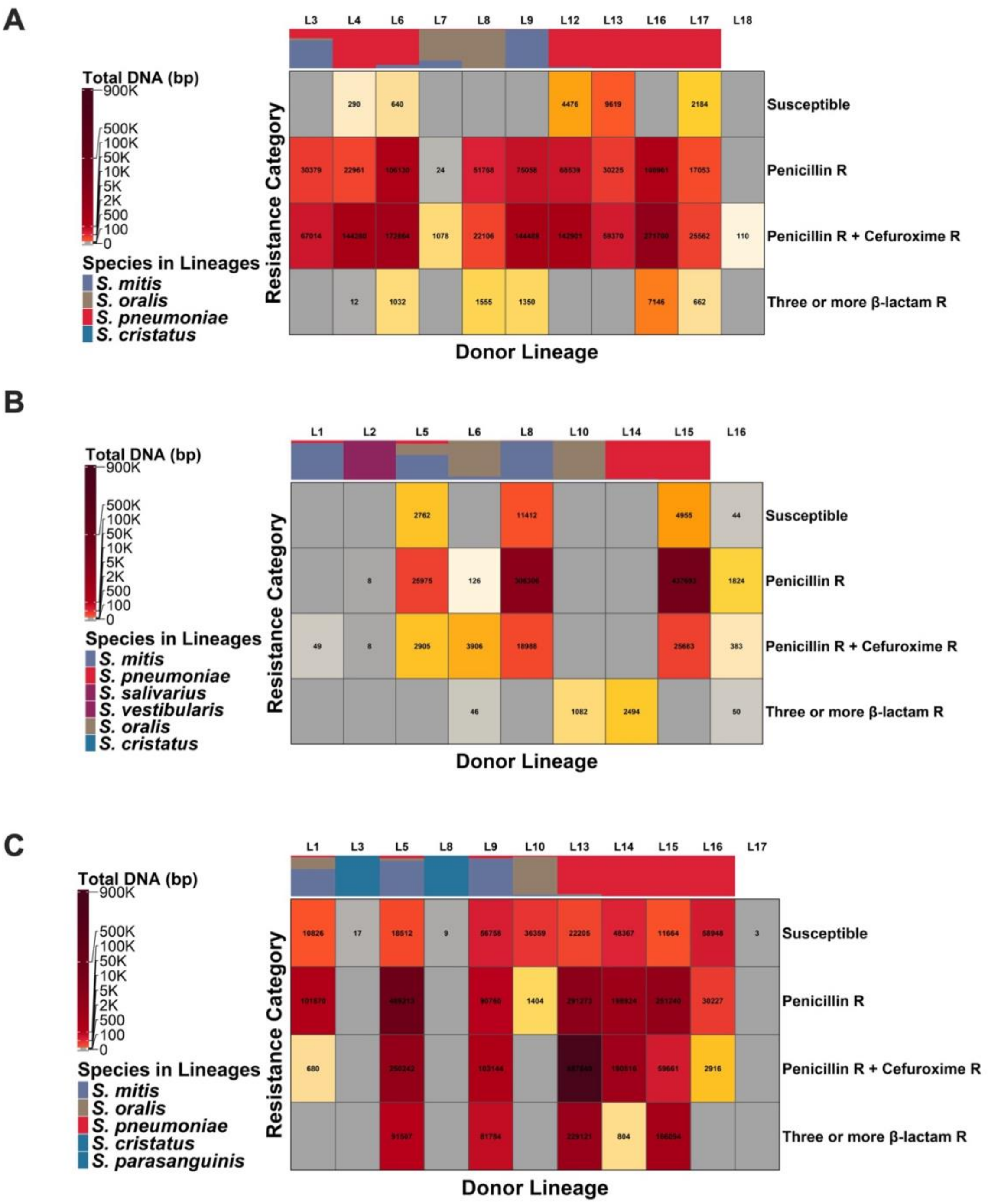


**Fig S4.** Recombination fragment profiles of penicillin binding protein genes across *S. pneumoniae* resistance categories for **(A)** *pbp1a*, **(B)** *pbp2b*, **(C)** *pbp2x*. Heatmaps of total recombination fragment length transferred from donor lineages to *S. pneumoniae* recipient resistance categories. Columns represent donor lineages ordered by species composition shown in the top color bar. Rows represent *S. pneumoniae* resistance categories according to minimum inhibitory concentration prediction. Each cell displays the total length of recombination fragments in base pairs. Color intensity shows total transferred DNA.

Seemann T. https://github.com/tseemann/abricate. Abricate. Github.
